# Transmission network for foot-and-mouth disease epidemic in Mar Chiquita - Argentina: A spatial modeling analysis

**DOI:** 10.1101/2025.09.26.678789

**Authors:** Salihu S. Musa, Laura Camila Lozano, Andrea Marcos, Guido A. König, Andrés Colubri

## Abstract

Foot-and-mouth disease (FMD) is a highly contagious viral disease with major implications for livestock production, food security, and economic stability. In this study, we investigate the transmission dynamics of the 2001 FMD outbreak in Mar Chiquita, Argentina, using a spatially explicit modeling framework. Transmission kernels based on inter-farm Euclidean distances are developed to estimate the probability of between-farm transmission and to reconstruct the likely transmission network. Several kernel formulations are evaluated, and parameters are estimated using maximum likelihood estimation techniques. We further assess the influence of isolation interventions by incorporating time-dependent kernel modifications that reflect reduced transmission risk following farm visits. Local transmission potential is quantified through farm-specific reproduction numbers, and transmission pathways are inferred using Markov Chain Monte Carlo (MCMC) simulations. Our results show that controlling geographic proximity alone does not reliably mitigate transmission risk and that delayed intervention can sustain epidemic spread. The reconstructed transmission network reveals spatial clusters of infections and temporal patterns consistent with field observations. These findings highlight the importance of timely isolation, active surveillance, and spatial risk assessment for effective outbreak control.

**Highlights:**

- Spatial transmission modeling captures FMD spread across 65 farms in Mar Chiquita, Argentina.
- Distance-based kernel functions estimated from epidemic data quantify between-farm infection risk.
- MCMC simulations reconstruct high-resolution farm-to-farm transmission networks.
- Intervention strategies (e.g., isolation) effectively reduce transmission probability and local R_0_.

## 1 Introduction

Foot-and-mouth disease (FMD) is a highly contagious viral disease that affects cloven-hoofed animals, such as cattle, pigs, sheep, goats, and deer [1–9]. FMD is caused by the Foot-and-Mouth Disease Virus (FMDV), a member of the picornavirus family [2, 4, 10]. The virus has seven serotypes, each of which can have many subtypes, making it challenging to develop long-lasting immunity. Contact with infected animals or contaminated objects like machinery, vehicles, or clothing contributes to the spread of FMD [2, 8]. Airborne transmission is also possible, particularly in areas with abundant animal life and within short distances [11, 12]. Modeling and experimental studies [13, 14] suggest that indirect transmissions via transport vehicles, cattle sheds, and farm visitors play a significant role in the farm-to-farm transmission pathway, beyond animal movements and other local transmission events. Fever, blisters or sores on the tongue, lips, gums, nostrils, and the area between the hooves are some of the symptoms of FMD [1, 2, 8, 15]. Lameness, loss of appetite, decreased milk output, and weight loss may occur in infected animals. The disease is rarely lethal to adult animals [2, 3, 16]. Nonetheless, it could cause significant economic losses due to diminished production, limits on animal mobility, and international trade bans on cattle and animal products.

Animals who recover from FMD infection might become carriers (a state in which the disease is not harmful to affected animals but can transmit to others) albeit this is more common in ruminant species [2, 17]. As the viral pathogens in infected animals are low and unstable, repeated oropharyngeal fluid tests may be required to detect potential carriers. Real-time RT-PCR tests are routinely employed in laboratories to test exposed animals; these assays are often divided into two groups, each targeting various sections of the RNA genome [2]. It is critical to identify previously infected animals from those who have been vaccinated against FMD. This is because a substantial proportion (up to 50%) of FMDV-infected animals might become carriers [18]. Carrier animals are clinically normal and can maintain this state for a long period (up to 2–3 years in cattle); those animals (such as African buffalo) could be a potential source of infection for other susceptible animals [2, 19]. However, more experimental studies are needed to demonstrate this on other livestock. The perceived risk presented by carriers has also significantly impacted the protections in place to manage the risks involved with international cattle movements. As a result, complete embargo, quarantine, and testing may be effective control tactics.

Transmission network modeling is an important area of analysis for FMD transmission - it integrates genomic and epidemiological data to infer “who infected whom” during disease outbreaks. Previous studies [20–22] examined the genetic diversity and molecular epidemiology of FMDV in Argentina, providing critical insights into the virus’s evolution and transmission dynamics during the 2000–2002 outbreaks. Several models have been developed to reconstruct transmission networks and estimate key parameters. For instance, Lau’s systematic Bayesian inference framework, implemented in the R package BORIS, has been successfully applied to the 2010 FMD outbreak in Japan; the resulting estimates have been found to improve overall accuracy when inferring transmission networks and timing of exposures [23]. Another study investigated the impact of altering the contact structure of the livestock movement network on disease dynamics. Rewiring the network through altered movement restrictions and redirection of movements between premises resulted in networks with higher clustering coefficients and lower network density, leading to smaller outbreaks [24]. Additionally, network analysis based on road networks and administrative districts has been used to analyze the spread of FMD and identify problems with the current method of setting preventive zones against the disease [25]. FMD has significant implications for agriculture, as it can spread rapidly and cause large outbreaks. Controlling FMD requires timely and effective intervention measures, including movement restrictions on affected animals, isolation, and culling could be used to prevent further spread of the disease [26]. Vaccines are available and most common in FMD-endemic regions but provide limited cross-protection against different strains [16, 27]. Furthermore, stringent biosecurity measures such as disinfection standards and surveillance programs are crucial in preventing and managing FMD transmission. Monitoring systems to monitor FMD outbreaks and emergency response plans to contain and eliminate the disease might be enacted during an epidemic. Although FMD does not pose a serious risk to human health, however, the impact of FMD on agricultural activities could be severe [28]. It can cause serious damage to the economy due to trade restrictions, livestock losses, and the cost of disease control methods. As a result, prevention, early detection, and quick treatment are imperative in reducing the impact of FMD on animal welfare and the global food supply chain.

Argentina reported its first FMD outbreak in 1886 through imported cattle. Subsequently, the outbreak (including the devastating 2001 epidemic) led to several outbreaks in subsequent years owing to the lack of effective control measures [8, 20, 21, 29–31]. Argentina has been constantly implementing an FMD vaccination campaign in some regions of the country to preserve its FMD-free status [29]. The risk of FMDV transmission into FMD-free zones through legal trading was assessed using quantitative risk analysis in [32], where intervention measures such as serological testing can be strictly imposed. Moreover, Brito et al. [22] examined the genesis and propagation of a substantial FMD outbreak in Argentina from 2000 to 2002, revealing insights into the virus’s genetic diversity and transmission dynamics. Furthermore, the efficiency of systematic mass vaccination programs against FMD in Argentina has been assessed by [33], with districts showing varying levels of protection.

The transmission dynamics of the FMD outbreak has been studied extensively in the literature in several settings (see, for instance, [2, 5, 6, 9, 11, 15, 34–36] and the references therein), ranging from disease epidemiology [2, 17, 37, 38], virus characterization [37], heterogeneity [10], diagnosis and surveillance [39], vaccination and other intervention measures [3–5, 16, 27, 34, 40], effect of temperature [41] as well as socio-economic impacts [42]. Although previous phylodynamic analyses have been completed [31], spatial analyses have not been explored for the Mar Chiquita outbreak. Furthermore, several prior works [23, 24, 43–45] have extensively used spatial epidemiological modeling to analyze animal disease transmission. Spatial analysis can help identify the source of virus exposure and any environmental factors that could enhance the virus transmission, which, in turn, helps control the spread of the virus [45]. Spatial analysis can be combined with epidemiological methods to evaluate potential spatial risk factors for FMD transmission [45, 46]. In particular, Lau et al. [43] proposed a Bayesian inference-based paradigm that takes into account epidemiological-evolutionary dynamics to explore the spread of FMD transmission dynamics. They discovered that their method could infer accurate epidemiological-evolutionary dynamics even when pathogen sequences and epidemiological data are incomplete or genetic sequences are only available for a fraction of exposures. This novel approach was extended by Firestone et al. [23] to integrate other characteristics and tested it on simulated and actual outbreak data from Japan. According to the authors’ findings, the modified model improved accuracy and could assess who infected whom via network reconstruction [23, 24], as well as provide the best estimate for crucial parameters such as transmissibility and infer critical linkages between clusters and features of farms involved in the outbreak [23]. While previous studies—such as [31]—have attempted to explore the phylodynamics of FMD outbreaks in Argentina, the spatial transmission patterns of the disease, particularly in regions like Mar Chiquita, remain underexplored. In this context, the development of a spatially explicit FMD transmission model becomes crucial for enhancing epidemic preparedness and informing targeted intervention strategies in the face of potential future outbreaks.

This study adopts a spatial modeling framework to investigate the 2001 foot-and-mouth disease (FMD) outbreak in Mar Chiquita, Argentina. Our models converge on a common infection timeline by mid-June [31], consistent with previous phylodynamic findings for the region. We formulate and compare multiple distance-based transmission kernels to estimate the probability of inter-farm transmission based on Euclidean distances. Using these kernel estimates, we reconstruct the most likely transmission network to infer “who infected whom” during the epidemic. The model also incorporates relevant epidemiological quantities to estimate infection rates and evaluate the impact of control interventions—particularly isolation measures—on the overall transmission dynamics. Furthermore, we compute the local reproduction number to assess spatial heterogeneity in epidemic risk across farms. Numerical simulations are presented to support analytic results, with implications for designing data-informed and timely intervention strategies in future FMD outbreaks. This work was supported in part by funding from the National Scientific and Technical Research Council (Consejo Nacional de Investigaciones Científicas y Técnicas, CONICET) in Argentina, with Laura Lozano as a Doctoral fellow.

## 2 Material and methods

### 2.1 Data

This study utilizes epidemiological data from the 2001 foot-and-mouth disease (FMD) outbreak in Mar Chiquita, Argentina, provided by the National Food Safety and Quality Service (SENASA) [47]. The dataset includes information from 65 livestock farms (denoted A1–A65) distributed across the region, with Farm A1 identified as the likely index case. The data encompass spatial coordinates, livestock composition, and daily case counts before and after intervention measures were implemented. Each farm recorded the total number of cattle, sheep, goats, and swine. However, for modeling purposes, our analysis focused exclusively on cattle and sheep, which together accounted for over 98% of the infected animals during the epidemic. Goats and swine were excluded due to their negligible contribution to overall transmission. The dataset includes detailed case counts of susceptible and infected animals prior to and following veterinary intervention (i.e., farm visits for isolation or culling). Specifically, the pre-intervention phase documented 74,006 healthy and 5,610 infected animals, while the post-intervention phase reported 81,705 healthy and 11,163 infected animals. This apparent increase in total animal counts post-intervention does not indicate a true rise in livestock populations. Instead, it reflects enhanced case reporting and census correction during the intervention period, where additional animals and infections were recorded that were either previously unreported or detected after follow-up investigations (due to additional cases reported). Such reporting artifacts are common in the context of field outbreaks, especially when surveillance efforts intensify following initial control measures. These case distributions are illustrated in Figure 1, where panels (a) and (b) display susceptible and infected animals before intervention, and panels (c) and (d) show the respective counts after intervention. The geographical distribution of the 65 farms is shown in Figure 2, where each point corresponds to a farm location within Mar Chiquita. The inter-farm distances were computed using the Haversine formula based on geographic coordinates (latitude and longitude), yielding a comprehensive distance matrix visualized in Figure 3. For data confidentiality and ownership reasons, precise farm coordinates are not publicly disclosed in this study. Any request for access to exact farm locations should be directed to the original data custodians, including Andrea and the National Food Safety and Quality Service (Servicio Nacional de Sanidad y Calidad Agroalimentaria, SENASA), who maintain the full dataset. All data utilized in this study were obtained from public sites and from the National Directorate of Animal Health at SENASA (farm information is available under request). Furthermore, the inclusion of distancebased kernel structures in the modeling framework is supported by established FMD transmission dynamics, where spatial proximity plays a critical role in farm-to-farm spread. The resulting kernel captures the spatial decay in transmission probability and is informed by the computed inter-farm distances.

**Figure 1:**
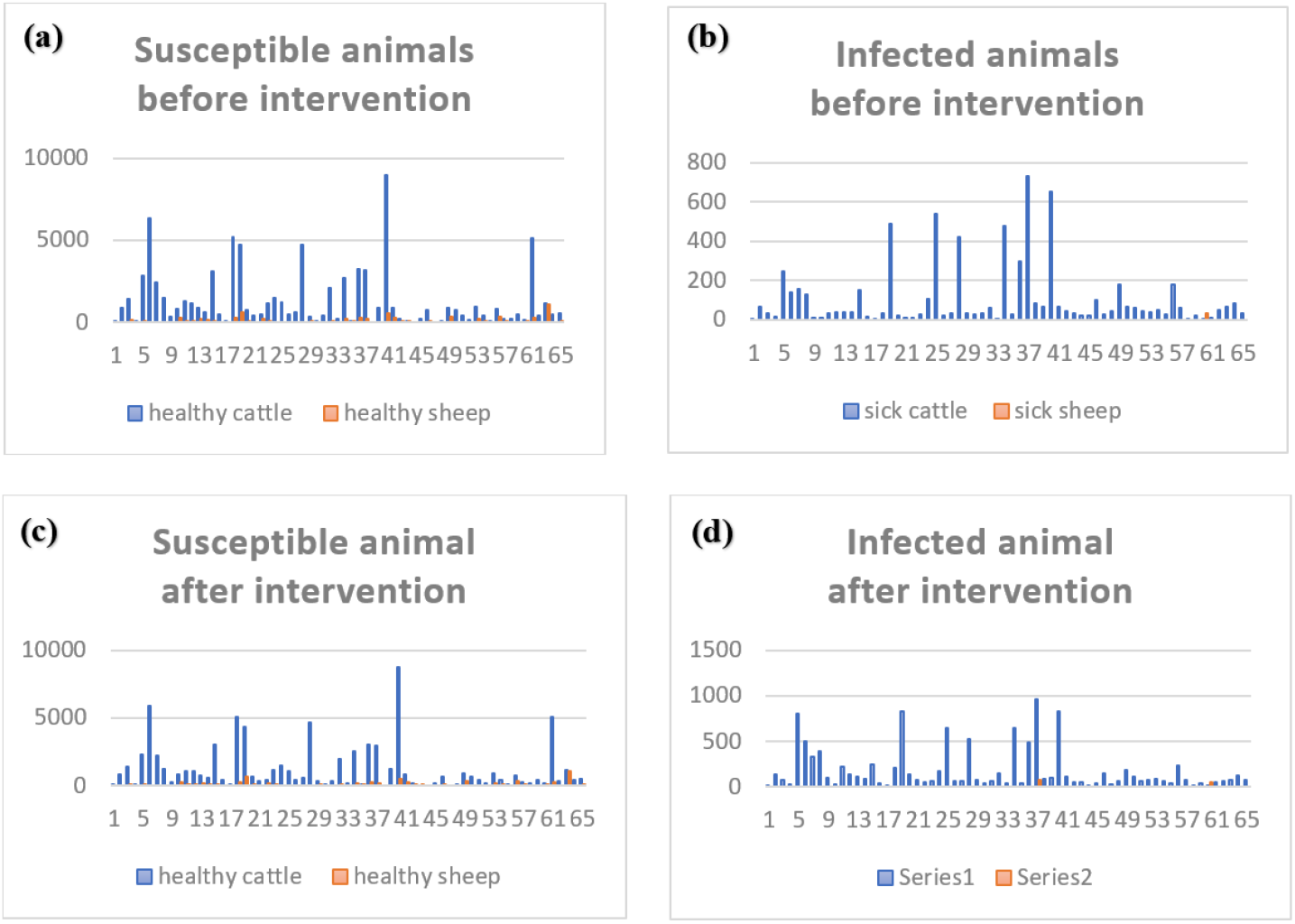
FMD cases reported in Mar Chiquita, Argentina (2001), across 65 farms. Panels (a) and (c) show the number of susceptible animals before and after the implementation of control measures, respectively, while panels (b) and (d) show the corresponding infected animal counts. In all panels, the X-axis represents farm IDs (A1–A65), and the Y-axis indicates the number of animals (susceptible or infected).

**Figure 2:**
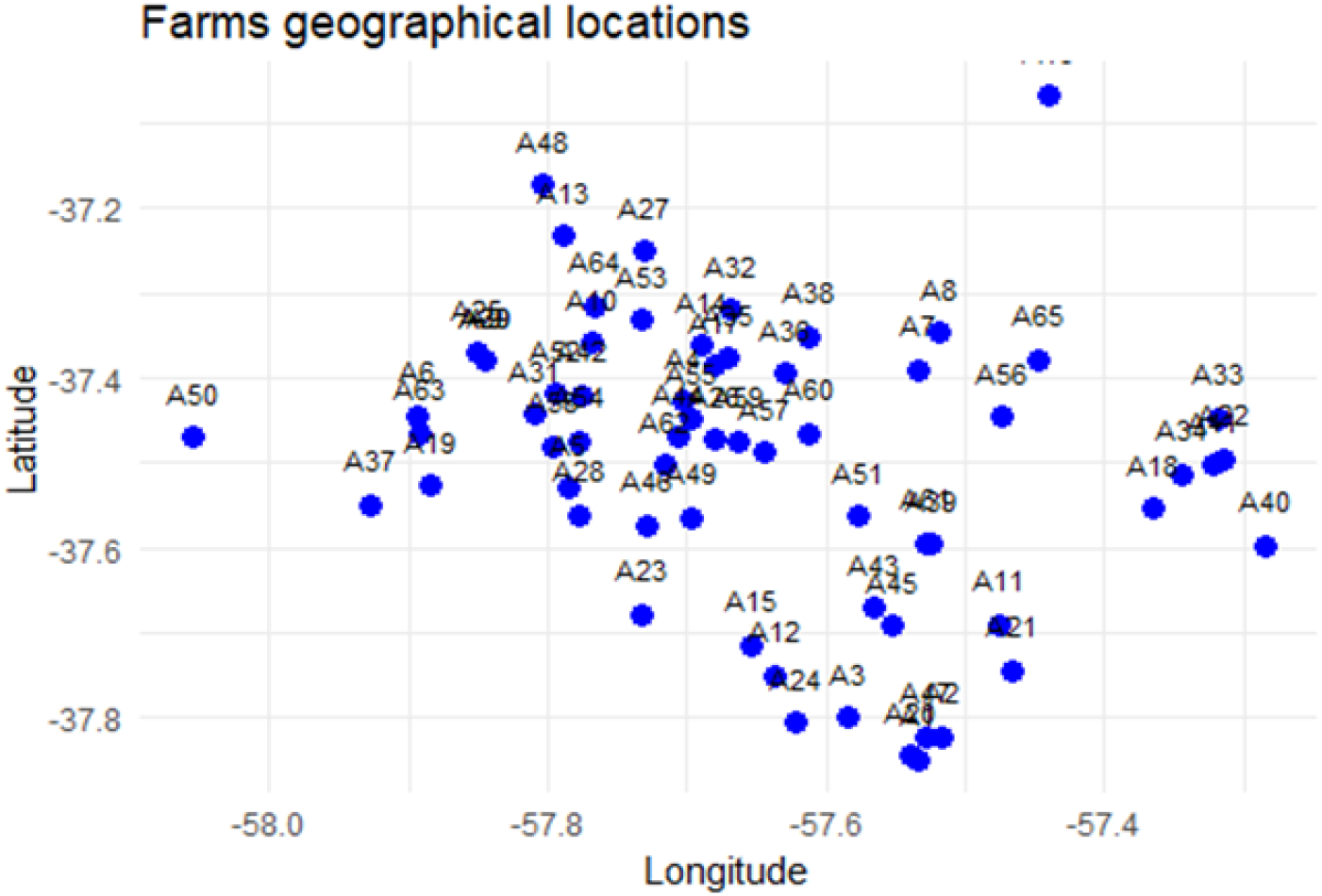
Spatial distribution of the 65 livestock farms in Mar Chiquita. Blue dots represent farm locations labeled A1 to A65.

**Figure 3:**
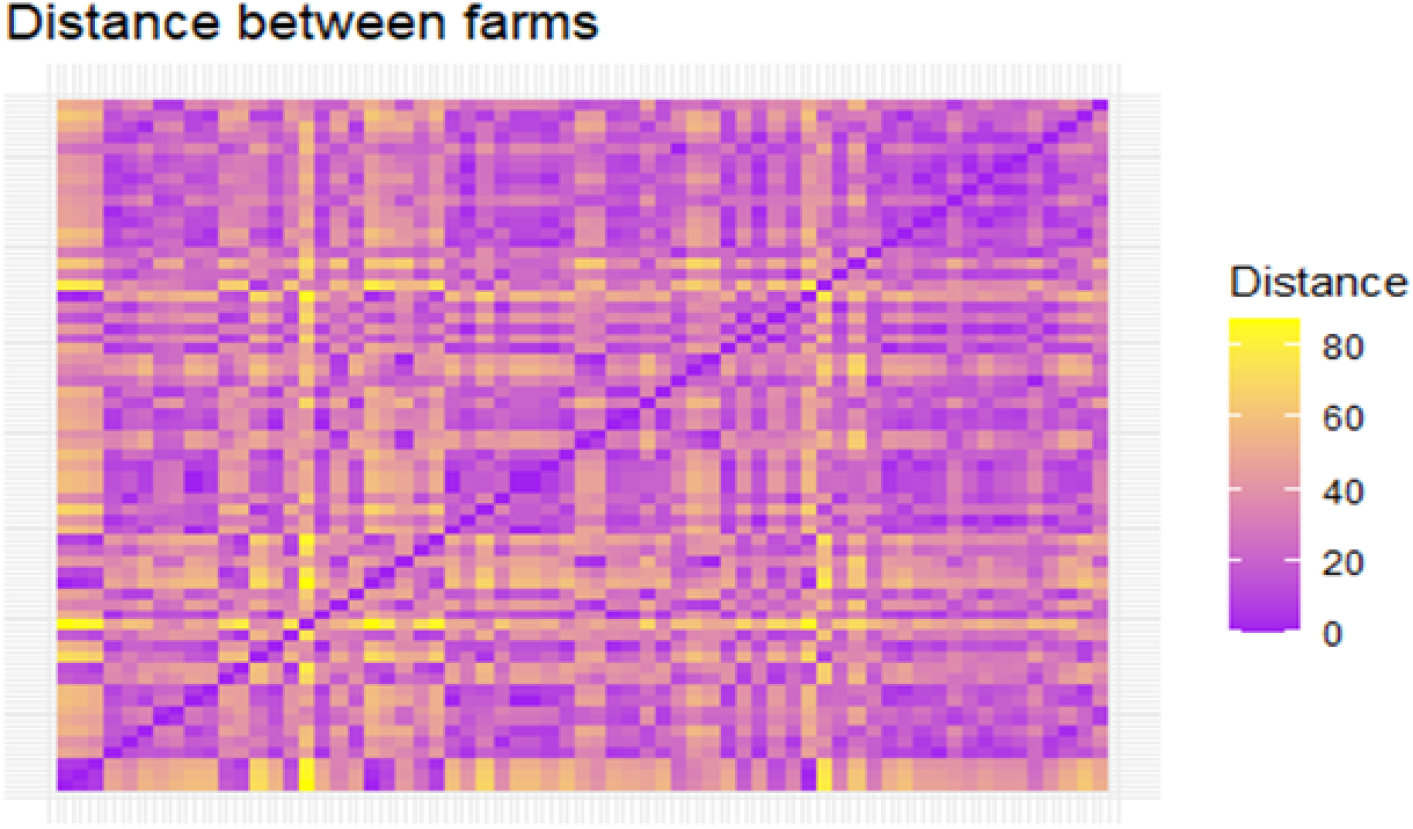
Inter-farm distance matrix for the 65 farms in Mar Chiquita. Color gradient indicates pairwise geographical distance, with purple denoting shorter and yellow longer distances. Distance is in kilometers.

### 2.2 Transmission kernel

During FMD epidemics, several transmission routes contribute to between-farm spread, including direct contact through animal movements, indirect contact via contaminated materials (e.g., vehicles, equipment, or human clothing), and airborne transmission. Incorporating all these routes into a dynamic model can be complex. However, spatial modeling provides a framework to estimate key epidemiological quantities—such as transmission rates, infection probabilities, contact structures, and reproduction numbers—that collectively describe the spread of infection [15, 26, 27, 35, 48–51]. The spatial transmission risk between an infectious farm *i* and a susceptible farm *j* at time *t* is modeled using a transmission kernel *h*(*d*_*ij*_, *t*), where *d*_*ij*_ denotes the Euclidean distance between the two farms. Following previous studies [15, 27, 35], we evaluate several forms of *h*(*d*_*ij*_, *t*), incorporating the effects of isolation measures and control strategies. Parameters for the kernel functions are estimated using maximum likelihood estimation (MLE) methods. These kernels allow us to model transmission probability as a decreasing function of distance. This approach has been used in various animal diseases such as avian influenza and classical swine fever [6, 7, 16, 34, 35, 52–54]. In this study, we apply these kernel-based approaches to estimate inter-farm transmission probabilities during the 2001 FMD outbreak in Mar Chiquita, Argentina.

To incorporate appropriate transmission kernels into the FMD model, we adopt the functional forms evaluated in [15, 34, 35, 55], particularly the three-parameter logistic expression:

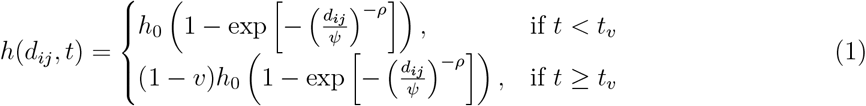

Here, *h*_0_ is the maximum transmission value (achieved when *d*_*ij*_ = 0), *ψ* is the kernel offset, and *ρ* is the kernel power shaping the spatial decay. The term *v* ∈ (0, 1) represents the effectiveness of isolation measures introduced at time *t*_*v*_. This logistic kernel is monotonically decreasing with distance and widely used for spatially structured epidemics. For example, Savill et al. [53] showed that Euclidean distance was a strong predictor of transmission risk during the FMD outbreak in Great Britain, except where geographical barriers were prominent.

To test robustness, we also consider alternative kernel formulations frequently used in the literature [15, 34, 35, 55]:

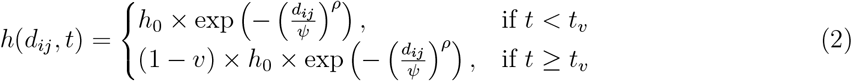

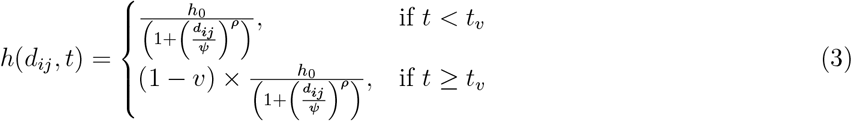

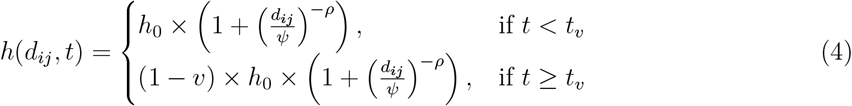

These kernel functions were simulated to compare their shapes using a common parameter set (*h*_0_ = 0.1, *ψ* = 1, *ρ* = 2), and the results are illustrated in Figure 4. The plot shows how each kernel decays with distance under pre-intervention conditions, highlighting differences in tail behavior and sensitivity to proximity.

**Figure 4:**
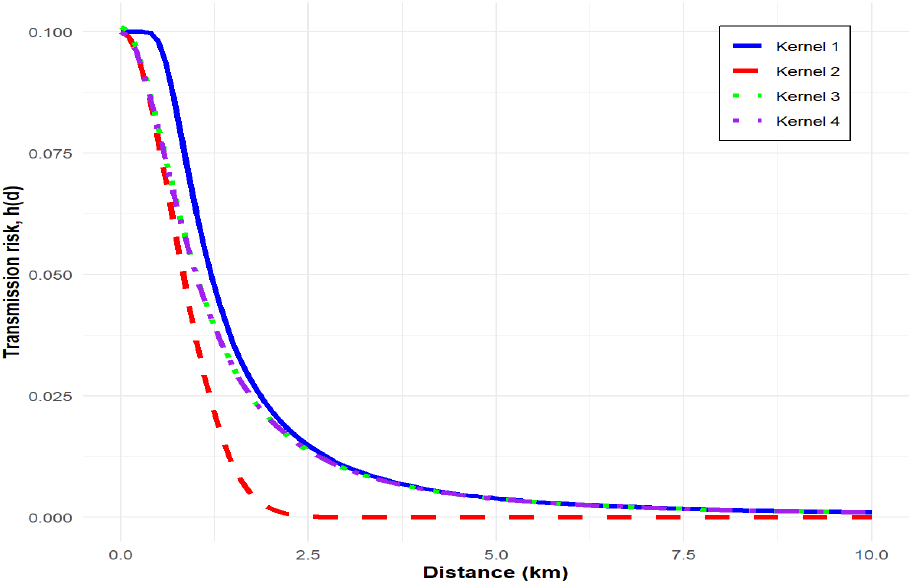
Spatial transmission kernels *h*(*d*) as a function of inter-farm distance *d*, comparing four functional forms used to model FMD spread in Mar Chiquita. All kernels represent pre-intervention transmission with parameters: *h*_0_ = 0.1, *ψ* = 1, and *ρ* = 2.

The following assumptions underlie the baseline transmission kernel model applied to the Mar Chiquita FMD outbreak:

i) *h* = 0 implies no infection occurs between farms during the epidemic period.
ii) If *d*_*ij*_ = 0, then the kernel reaches its maximum value *h*_0_.
iii) For *d*_*ij*_ 0 and *t < t*_*v*_, transmission follows *h*(*d*_*ij*_, *t*), reflecting no intervention.
iv) For *d*_*ij*_≠ 0 and *t* ≥ *t*_*v*_, transmission is reduced by isolation: *h*(*d*_*ij*_, *t*) = (1 −*v*) ×*h*(*d*_*ij*_, *t*). When *v* → 0, intervention is minimal; when *v* → 1, intervention is complete.

These kernel models were fitted to outbreak data via maximum likelihood estimation to identify the most appropriate formulation for capturing the observed epidemic patterns in Mar Chiquita.

### 2.3 FMD spatial epidemic model

The spatial model employed in this study builds upon the foundational framework developed by Keeling et al. [6], which was subsequently extended and applied in numerous FMD modeling studies [15, 26, 27, 35, 50, 51]. We adapted this spatially explicit model to examine the 2001 FMD epidemic in Mar Chiquita, Argentina, with the goal of quantifying the transmission dynamics between farms and evaluating the impact of intervention strategies such as isolation. In this model, the force of infection from an infectious farm *i* to a susceptible farm *j* at time *t* is determined by the number of animals on both farms, their species-specific susceptibility and transmissibility, and the inter-farm distance. Specifically, the transmission rate is defined as:

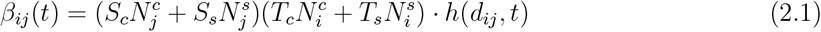

Here, 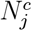 and 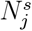 represent the number of susceptible cattle and sheep on farm *j*, respectively, while 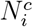 and 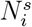 denote the number of infectious cattle and sheep on farm *i*. The parameters *S*_*c*_ and *S*_*s*_ denote the per capita susceptibility of cattle and sheep, respectively, and *T*_*c*_ and *T*_*s*_ represent their corresponding transmissibility. The term *h*(*d*_*ij*_, *t*) is the spatial transmission kernel that captures the decay of infection risk as a function of Euclidean distance *d*_*ij*_ between farms *i* and *j*, possibly modulated by time-dependent intervention effects such as movement restrictions or isolation. This formulation captures all four species-specific transmission pathways: cattle-to-cattle (*S*_*c*_*T*_*c*_), sheep-to-cattle (*S*_*c*_*T*_*s*_), cattle-to-sheep (*S*_*s*_*T*_*c*_), and sheep-to-sheep (*S*_*s*_*T*_*s*_), as also considered in earlier modeling frameworks [6, 26, 35, 50]. When the kernel is assumed distance-independent (i.e., uniform contact), the model reduces to a mass-action form [35]. However, by incorporating spatial heterogeneity via *h*(*d*_*ij*_, *t*), the model accounts for more realistic local transmission dynamics.

The probability that farm *j* becomes infected on day *t* is then given by:

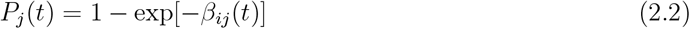

This expression assumes that infections follow a Poisson process and that transmission events are independent across farm pairs. The daily infection probabilities across all farms are aggregated to form a likelihood function for parameter estimation. These probabilities are especially important in likelihood-based inference schemes, including maximum likelihood estimation (MLE) and Markov chain Monte Carlo (MCMC), which are commonly used to estimate transmission kernel parameters [15, 35].

Although the species-specific susceptibility and transmissibility parameters may not reflect absolute biological values, they provide an interpretable and consistent way to capture relative transmission risk, especially in data-limited settings. Importantly, previous studies [6, 10, 15] have shown that human-mediated contact, such as via vehicles, workers, or shared equipment, can modulate effective transmission between farms, further motivating the use of distance-dependent transmission kernels. By combining heterogeneous host structure, spatial distance, and intervention dynamics, this spatial epidemic model enables the reconstruction of inter-farm transmission patterns and the evaluation of intervention strategies in the context of a real-world epidemic. The modeling choices are consistent with prior applications to FMD outbreaks in Japan [15], Great Britain [35], and avian influenza [52], thereby supporting both methodological rigor and policy relevance.

### 2.4 Parameter estimation

To estimate the parameters of the transmission kernel *h*(*d*_*ij*_, *t*), we employed a maximum likelihood estimation (MLE) approach based on the spatial epidemic framework. The kernel functions used—specified in Equations (1)–(4)—provide simplified yet analytically tractable representations of complex contact processes between infected and susceptible farms [6, 51]. In particular, the three-parameter formulation of Equation (1), which includes the kernel amplitude *h*_0_, the distance scaling parameter *ψ*, and the shape parameter *ρ*, was primarily utilized to fit the empirical data. This method follows prior works that applied MLE to epidemic data to identify kernel structure and parameter values [15, 35, 50, 52].

To construct the likelihood function, we define the force of infection on a susceptible farm *j* at time *t* as:

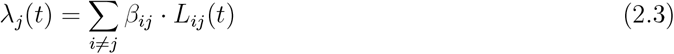

where the summation runs over all currently infectious farms *i*, and *L*_*ij*_(*t*) is an indicator function that takes the value:

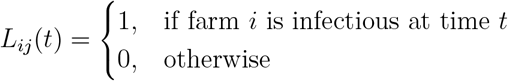

The probability that a susceptible farm *ϕ* becomes infected at time *t* is given by:

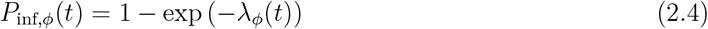

Meanwhile, the probability that farm *ϕ* remains uninfected up to day *t* is expressed as:

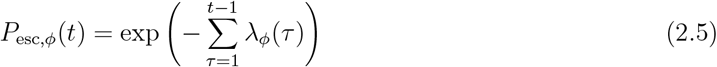

The overall likelihood function for the epidemic is then given by:

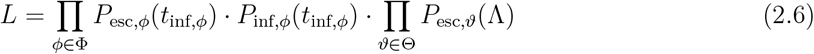

Here, Φ is the set of all farms that became infected during the outbreak, Θ is the set of farms that remained uninfected, and Λ is the total duration of the epidemic (from the first to the last reported infection). Taking the logarithm of the likelihood gives the log-likelihood function, which simplifies computation:

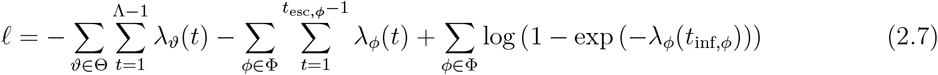

The kernel parameters *h*_0_, *ψ*, and *ρ* were estimated by numerically maximizing ℓ, separately for the pre-intervention phase (*v* = 0) and post-intervention phase (0 < *v* ≤ 1). These fitted parameters quantify how spatial distance influences transmission likelihood before and after control measures were implemented.

Model comparisons were also performed across all four kernel formulations (Equations 1–4) using their corresponding log-likelihood values. Among them, the formulation in Equation (1) provided the best fit to the observed outbreak data. The estimated parameter values and comparative performance are reported in Table 2, and simulated kernel profiles are visualized in Figure 4.

**Table 1:** Maximum likelihood estimation of the transmission kernel parameters.

| Kernel | transmission kernel ($h_0$) | | kernel offset ($\psi$) | | kernel power ($\rho$) | |
| --- | --- | --- | --- | --- | --- | --- |
| | $v = 0$ | $v = 0.5$ | $v = 0$ | $v = 0.5$ | $v = 0$ | $v = 0.5$ |
| 1 | 0.00498558 | 0.0007365943 | 0.734190956 | 0.0092264328 | 1.735751955 | 0.1252223836 |
| 2 | 0.1897069 |  | 3.5070398 |  | 2.0571439 |  |
| 3 | 2.805286e-02 |  | 3.403835e-16 |  | 1.770689e+01 |  |
| 4 | 8.381655e-01 |  | 7.653310e-06 |  | 4.585712e+01 |  |

**Table 2:** Log-likelihood estimates of the model performance for different transmission kernels.

| Kernel function | LL ($v = 0$) | LL ($v = 0.5$) |
| --- | --- | --- |
| 1 | -36775.87 | -265136.6 |
| 2 | -440880 |  |
| 3 | -2872850 |  |
| 4 | -2856799 |  |

**Table 3:** Summary of description and estimated values of model’s parameters.

| Parameter | Notation | Value ($v = 0$) | Value ($v = 0.5$) |
| --- | --- | --- | --- |
| Per capita cattle susceptibility | $S_c$ | 9.08640139 | 3.81057613 |
| Per capita sheep susceptibility | $S_s$ | 0.67409970 | 0.02049746 |
| Per capita cattle transmissibility | $T_c$ | 0.02959448 | 0.02431182 |
| Per capita sheep transmissibility | $T_s$ | 0.05095511 | 0.05455361 |

### 2.5 Construction of transmission network

Transmission network models are widely employed in epidemiological research to reconstruct likely infection pathways—i.e., to infer “who infected whom”—during outbreaks [23, 24]. In this study, we utilized spatially derived transmission probabilities to reconstruct the farm-to-farm transmission network for the 2001 FMD outbreak in Mar Chiquita, Argentina. These reconstructed pathways allow identification of key source farms, transmission clusters, and directional spread patterns, which can help guide targeted interventions to effectively contain future outbreaks.

We implemented a modified network reconstruction approach inspired by Sellman et al. (2018) and Firestone et al. (2015), which infers likely transmission trees using spatial transmission probabilities and temporal constraints. As shown in Figure 7, the inferred transmission network was constructed in a random spatial environment, accounting for the estimated probability of infection between each pair of farms. Networks generated under both the original and the normalized transmission models are provided in Figure 7, highlighting key differences and validating the robustness of the reconstructed pathways. The root node (initial infection source) was inferred with strong model support, corresponding to the index case recorded on the earliest outbreak date. The sequence of secondary and tertiary transmissions in the network followed spatial patterns consistent with both the estimated kernel structure and the chronological order of reported clinical signs. To provide a comparative baseline, we also applied the pairwise algorithm described in [23], which simulates direct farm-to-farm transmission events based solely on infection probabilities calculated using the spatial model. Each infectious–susceptible farm pair is treated independently, and the infection event is evaluated as a Bernoulli process. While conceptually simple and easy to implement, the pairwise method becomes computationally intensive for large populations, particularly when transmission probabilities must be calculated dynamically or when full pairwise distances are required.

#### Pairwise algorithm

The simplified pseudocode for the pairwise transmission simulation is as follows:

~~~
Probability P_ij as defined by Eq (12).
for each infectious node i {
   for each susceptible node j {
       r = uniform random number (0, 1)
      if r < P_ij { j is infected }
   }
}
~~~

While useful for comparative purposes, the pairwise algorithm was not used to generate the final transmission tree due to its computational limitations and lack of integration with temporal progression. Instead, our final reconstruction relied on a likelihood-informed MCMC approach described in Section 2.5.

### 2.6 MCMC simulations framework

To reconstruct the most probable transmission tree of the FMD outbreak in Mar Chiquita, we applied a Markov Chain Monte Carlo (MCMC) approach grounded on transmission probabilities computed from Equation (A1) after calibrating the model parameters via maximum likelihood estimation. Farms that were not infected during the outbreak were assigned a fixed transmission probability of zero, while those observed to have cases were attributed probabilistic infection events based on the estimated transmission kernel. The MCMC algorithm commenced from a randomly generated network and iteratively modified infection histories by selecting a random time point uniformly from the outbreak duration and reassigning subsequent infections according to the transition probability matrix. Reassignment was constrained to align with the reported dates of clinical onset on each farm. The time from infection to isolation was modeled using a gamma distribution, with shape and rate parameters fitted from observed epidemiological data. We performed 25 independent runs of 10,000 iterations each, initialized with distinct random networks to explore a broad configuration space. The most probable network among these initial runs served as the starting point for a final extended simulation of 250,000 iterations. The network with the highest posterior probability from this final run was selected as the best-supported transmission tree.

## 3 Results

The 2001 FMD epidemic in Mar Chiquita, Argentina, involved 65 spatially distributed farms with varying herd compositions. Figures 1 and 2 summarize the number of susceptible and infected animals on each farm before and after intervention, and the geographic positions of all farms (A1–A65), respectively. The epidemic began on May 18 with the index case reported at farm A1 and ended by August 18. Across the outbreak, 5,610 infections occurred prior to intervention and 11,163 after. Farms comprised cattle, sheep, goats, and swine; however, consistent with our modeling assumptions (see Section 2), only cattle and sheep were included in the analysis, as goats and swine contributed minimally to the overall transmission dynamics during the outbreak. Figure 3 visualizes the inter-farm distances derived from geographic coordinates, ranging from 3.2 km (A1 to A2) to 78.9 km (A1 to A48). These distances informed the spatial kernel and probability of infection between farms.

To evaluate spatial dependence in transmission, we fitted four distance-based kernel functions to the outbreak data. Maximum likelihood estimation (MLE) was used to identify the best-fitting kernel model before and after intervention implementation. Figure 5 displays the transmission kernel matrix, where color intensity reflects infection potential: darker red indicates higher transmission. Loglikelihoods for kernel functions (1)–(4) were compared. Under intervention, kernel (1) yielded the best fit (LL = -265136.6), outperforming kernel (2) (LL = -440880), kernel (3) (LL = -2872850), and kernel (4)(LL = -2856799). Parameter estimates for kernel (1) under intervention were: *h*_0_ = 0.0007366, *ψ* = 0.0092264, and *ρ* = 0.1252. Without intervention, kernel (1) remained optimal (LL = -530273.2), with *h*_0_ = 0.004986, *ψ* = 0.07419, and *ρ* = 1.7358. Simulation of kernel functions (1)–(4) is shown in Figure 5, highlighting the spatial decay in transmission risk.

**Figure 5:**
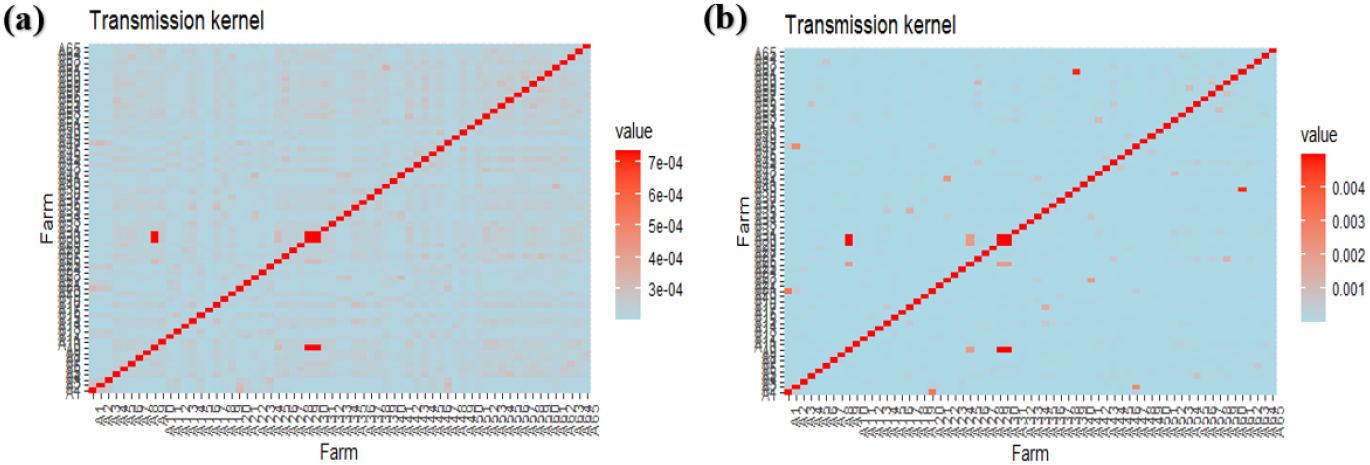
Estimated transmission kernel matrix for the 2001 FMD outbreak in Mar Chiquita. Panel corresponds to pre-intervention; panel (b) post-intervention. Red indicates higher transmission risk.

Using the best-fit kernel, we simulated infection probabilities across all farm pairs (Figure 6). Before intervention (panel a), several farm pairs exhibited high transmission probability (dark purple), which substantially declined following intervention (panel b, orange-to-yellow). These results support the hypothesis that timely interventions, such as movement restrictions and isolation, significantly reduce transmission likelihood between farms.

**Figure 6:**
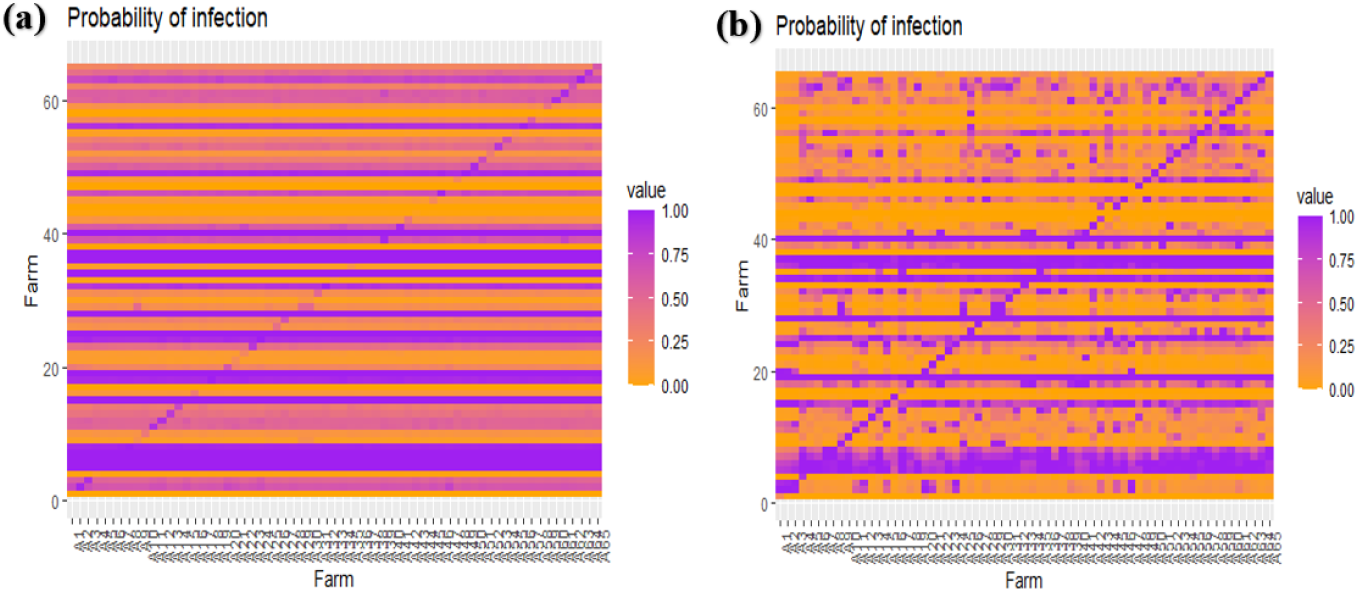
Simulated inter-farm infection probabilities. (a) Before intervention, (b) After intervention. Higher probabilities are shown in purple; lower in orange.

The spatially explicit model and estimated transmission probabilities were used to reconstruct the FMD transmission network using a modified pairwise transmission algorithm (Figure 7). Nodes represent farms, and directed edges represent inferred infection routes. Before intervention, dense interconnections between farms were observed, indicating high transmission risk. After intervention, the network became sparser, suggesting effective containment. This reconstruction helps identify high-risk farms and potential super-spreaders.

**Figure 7:**
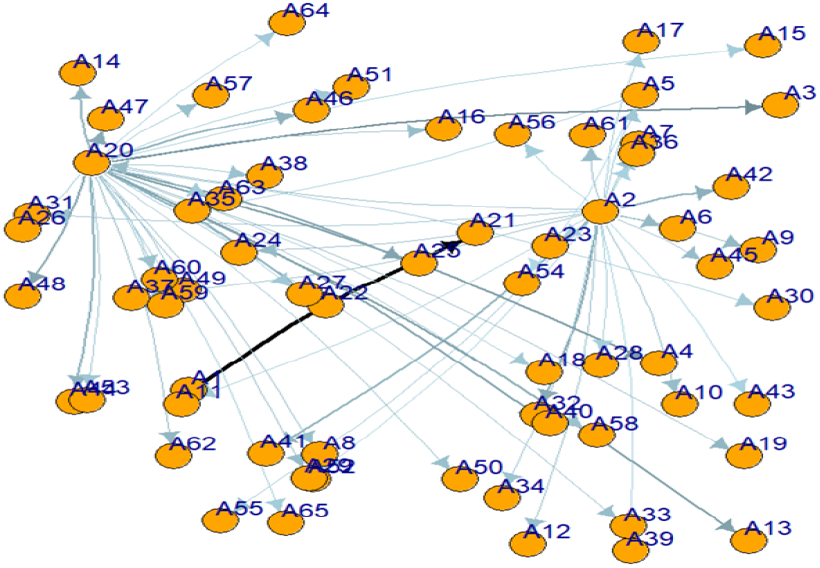
Inferred transmission network among 65 farms in Mar Chiquita based on kernel-estimated infection probabilities.

Figure 8 presents the most probable transmission tree reconstructed using Markov Chain Monte Carlo (MCMC) simulations. The network reveals a sequential spatial spread: initially emerging in the southeastern cluster at farm A1, spreading northwest by June 11, and reaching the northeastern farms by mid-June. This pattern aligns with prior phylogeographic evidence. Despite some timing discrepancies, both genetic and probabilistic reconstructions converged on similar spatial trajectories. Figure 8 presents the most probable transmission tree reconstructed using MCMC simulations. The network reveals a sequential spatial spread: initially emerging in the southeastern cluster at farm A1, spreading northwest by June 11, and reaching the northeastern farms by mid-June. This pattern aligns with prior phylogeographic evidence. Despite some timing discrepancies, both genetic and probabilistic reconstructions converged on similar spatial trajectories, and the MCMC-based network—derived from spatial kernel estimates and onset data—yielded a high-resolution reconstruction of farm-to-farm transmission pathways during the outbreak.

**Figure 8:**
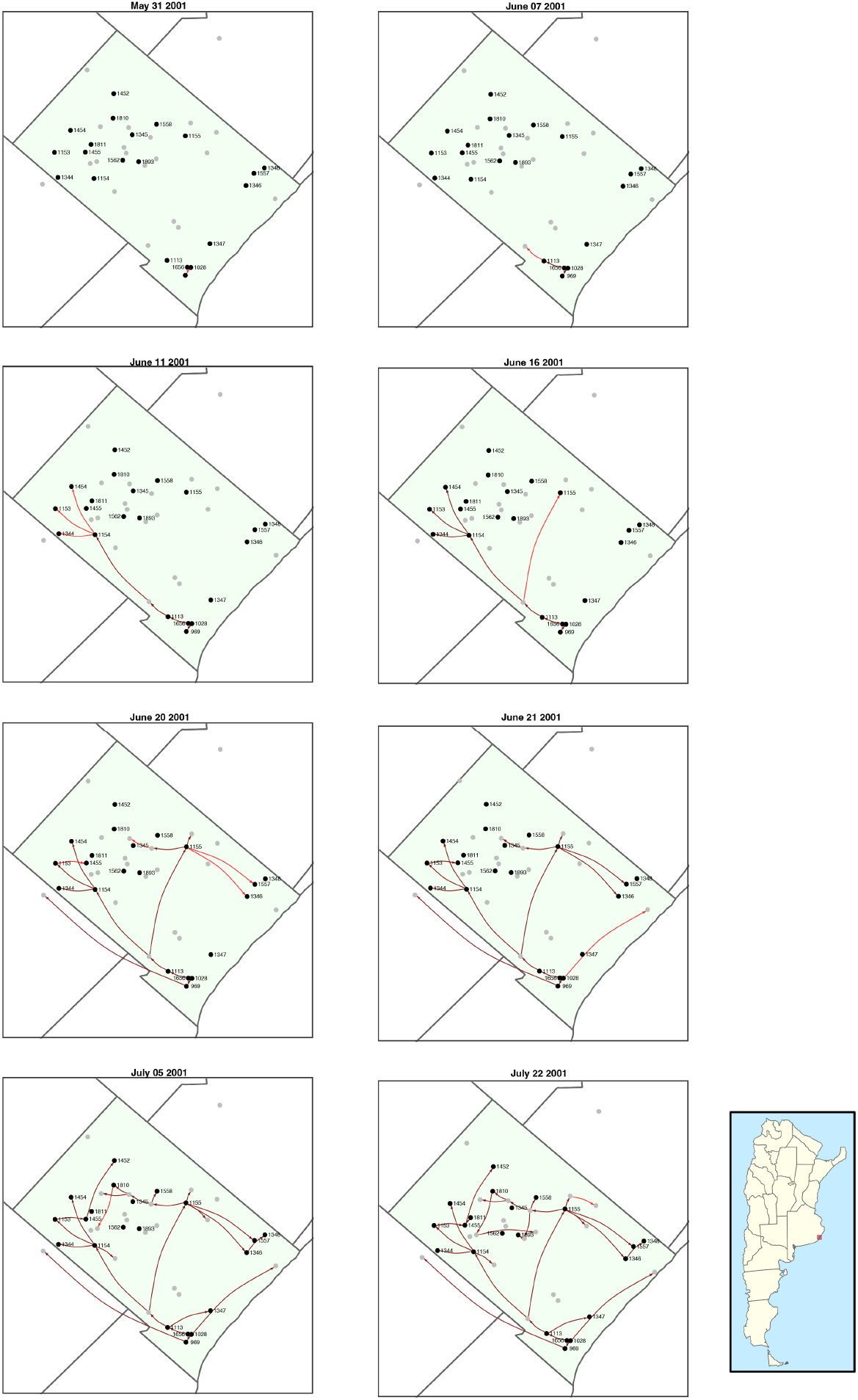
Reconstructed FMD transmission network among farms in Mar Chiquita during the 2001 epidemic. The network was inferred using Markov Chain Monte Carlo (MCMC) simulations based on estimated transmission probabilities between farms. Nodes represent farms, and directed edges indicate likely infection pathways. The color gradient reflects the estimated time of infection, with darker nodes infected earlier in the outbreak. The sequence of transmission shows initial spread from the southeastern farms, followed by infection of the northwestern cluster and later the northeastern farms. This pattern supports the hypothesis of spatially driven transmission and corresponds with observed epidemic data and phylogeographic evidence.

Although phylogenetic evidence places the northwestern infections slightly earlier (mid-May), both models converged on a similar infection timeline by mid-June, with the MCMC-based reconstruction—integrating kernel-based probabilities and onset data—revealing detailed transmission trajectories despite incomplete contact tracing.

## 4 Discussion

This study employed a spatial modeling framework to analyze the dynamics of foot-and-mouth disease (FMD) transmission during the 2001 outbreak in Mar Chiquita, Argentina. We developed and calibrated multiple distance-based transmission kernels to assess inter-farm transmission risk and investigate the effects of intervention measures, such as isolation. Our approach, which integrates spatial heterogeneity, maximum likelihood estimation, and probabilistic transmission modeling, reveals key insights into the epidemic spread and control.

We first estimated the transmission kernel based on inter-farm Euclidean distances, capturing the probability of infection from an infectious to a susceptible farm. Among the four candidate kernel functions examined, the best fit was obtained using Kernel 1—a modified exponential kernel with power transformation—as evidenced by the highest log-likelihood value (LL = -265136.6 with intervention; see Figure 5 and Table 2). The estimated kernel parameters before and after intervention (*h*_0_, *ψ*, and *ρ*) indicated a modest decline in transmission intensity following control measures. This relatively small change in parameter values may reflect continued movement of personnel, equipment, or vehicles between farms despite formal movement restrictions—a phenomenon previously reported in FMD outbreak investigations [15, 35, 56].

The corresponding transmission probabilities derived from these kernels (Figure 6) show that preintervention transmission likelihoods were concentrated near unity, whereas post-intervention probabilities dropped substantially. Although the estimated kernel parameters changed only modestly, the resulting transmission probabilities declined more sharply due to the nonlinear structure of the kernel function and its sensitivity to short-range transmission. This contrast underscores the importance of timely intervention strategies such as isolation and culling, as even moderate reductions in kernel parameters can produce significant epidemiological impact. Our analysis also confirmed that proximity alone does not fully explain transmission risk, suggesting that other factors—such as indirect contact pathways and farm-level heterogeneities—contribute meaningfully, consistent with findings from [35, 48, 50]. In line with our spatial modeling framework, the estimated local reproduction numbers (Figures A1 and A2, Supplementary Material) demonstrated that inter-farm transmission alone was insufficient to sustain widespread outbreaks, emphasizing the critical role of early intervention and non-distance-dependent transmission routes.

The reconstructed transmission network (Figure 7) offers critical insights into the farm-level spread of FMD in Mar Chiquita. Before the implementation of control measures, the network exhibited a dense web of connections, indicative of intense transmission potential across farms. Following intervention, the network structure became notably sparser, reflecting the reduction in transmission likelihood and highlighting the effectiveness of isolation and control strategies. This reconstruction not only corroborates the spatial dynamics of the epidemic but also pinpoints high-risk farms and potential super-spreaders, providing a valuable tool for prioritizing surveillance and targeted interventions in future outbreaks.

One of the significant contributions of this study is the use of Markov Chain Monte Carlo (MCMC) methods to reconstruct the transmission network with high spatial and temporal resolution. The inferred network (Figure 8) revealed a clear directional spread: beginning in the southeastern farms (notably A1), progressing northwest by June 11, and eventually reaching the northeastern sector by mid-June. Although the inferred timeline for northwest infections lagged behind phylogenetic estimates (mid-May), both approaches converged by mid-June, underscoring the robustness of integrating spatially explicit transmission kernels with probabilistic inference. The combined framework produced a detailed reconstruction of farm-to-farm transmission pathways, demonstrating its utility in settings where traditional contact-tracing data are limited. These findings are consistent with previous FMD transmission reconstructions, including those reported for the 2010 outbreak in Japan [57].

Our results affirm the utility of kernel-based spatial models in characterizing disease transmission patterns. The directional transmission structure observed supports the hypothesis of localized spread, possibly influenced by short-range movement of animals or shared infrastructure. The integration of kernel estimates with MCMC sampling enabled us to infer “who infected whom” with a high degree of temporal and spatial resolution. This approach provides an effective framework for outbreak reconstruction, especially in contexts with limited contact-tracing data, as supported by related works [6, 23, 24, 51, 56]. While the model captures key spatial features, some simplifications remain. The infectious period was treated uniformly across farms, and potential variability in host susceptibility and farm-level biosecurity was not explicitly incorporated. These limitations could be addressed in future extensions of the model by integrating temporal dynamics and farm-specific covariates [10, 41].

In conclusion, this study presents a comprehensive spatial modeling framework for analyzing the transmission dynamics of foot-and-mouth disease. Applying this framework to the 2001 Mar Chiquita outbreak enabled us to estimate transmission kernels, infer infection probabilities, and reconstruct detailed transmission pathways using MCMC simulations. Our findings emphasize the importance of spatial proximity, timely intervention, and high-resolution modeling in managing FMD epidemics. The methodology developed here holds promise for application in future outbreaks, facilitating rapid and informed decision-making, particularly in regions with dense livestock populations and constrained surveillance capacity [23, 24, 56, 57].

## Declarations

### Ethics approval and consent to participate

Not applicable.

### Data availability

All data utilized in this study were obtained from public sites and from the Dirección Nacional de Sanidad Animal, Servicio Nacional de Sanidad y Calidad Agroalimentaria (farm information is availiable under request).

### Code availability

The R scripts used in the analyses reported in this manuscript are available in the following GitHub repository: https://github.com/colabobio/fmd-mar-chiquita-spatial-model

## Acknowledgements

The authors thank the Dirección Nacional de Sanidad Animal, Servicio Nacional de Sanidad y Calidad Agroalimentaria providing access to the farm-level and outbreak data used in this study. Their support and collaboration were essential to the successful completion of this work..

## Authors’ Contributions

SSM performed the computational analyses, interpreted results, and wrote the manuscript, AC supervised the project, performed computational analyses, interpreted results, and revised the manuscript, KAG and LCL provided the data, interpreted results, and revised the manuscript, and AM authorized use of the data and provided feedback. All authors gave final consent for publication.

## Appendices

### A1 Basic reproduction number

The basic reproduction number (*R*_0_) is a fundamental quantity in infectious disease modeling that describes the expected number of secondary infections generated by a single infected unit in a fully susceptible population. In the context of farm-level transmission, *R*_0_ provides an estimate of the average number of farms that a single infected farm is expected to infect. It serves as a threshold indicator: if *R*_0_ *>* 1, the infection may spread and lead to an epidemic; if *R*_0_ *<* 1, the disease is likely to die out.

In this study, we used a spatially explicit formulation incorporating transmission kernels and stochastic infectious periods to estimate the local reproduction number for each farm involved in the 2001 FMD outbreak in Mar Chiquita, Argentina. This method is consistent with approaches applied in previous studies [34, 52]. The expression for the local reproduction number for farm *j* is given by:

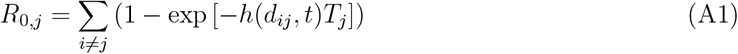

where *h*(*d*_*ij*_, *t*) is the transmission kernel as a function of the Euclidean distance between farms *i* and *j*, and *T*_*j*_ is the infectious period of farm *j*, drawn from a gamma distribution estimated from outbreak data.

Using the fitted transmission kernel (Figure 5), we computed local *R*_0_ values for each farm to assess early-stage transmission potential. Figures A1 and A2 display the resulting reproduction numbers before and after implementation of control measures. The analysis suggests that inter-farm transmission alone was unlikely to generate large outbreaks, highlighting the greater risk posed by within-farm and indirect transmission routes, particularly via human and vehicle movement. These findings underscore the importance of timely intervention, especially in preventing cross-farm spread during the early phase of the outbreak.

**Figure A1:**
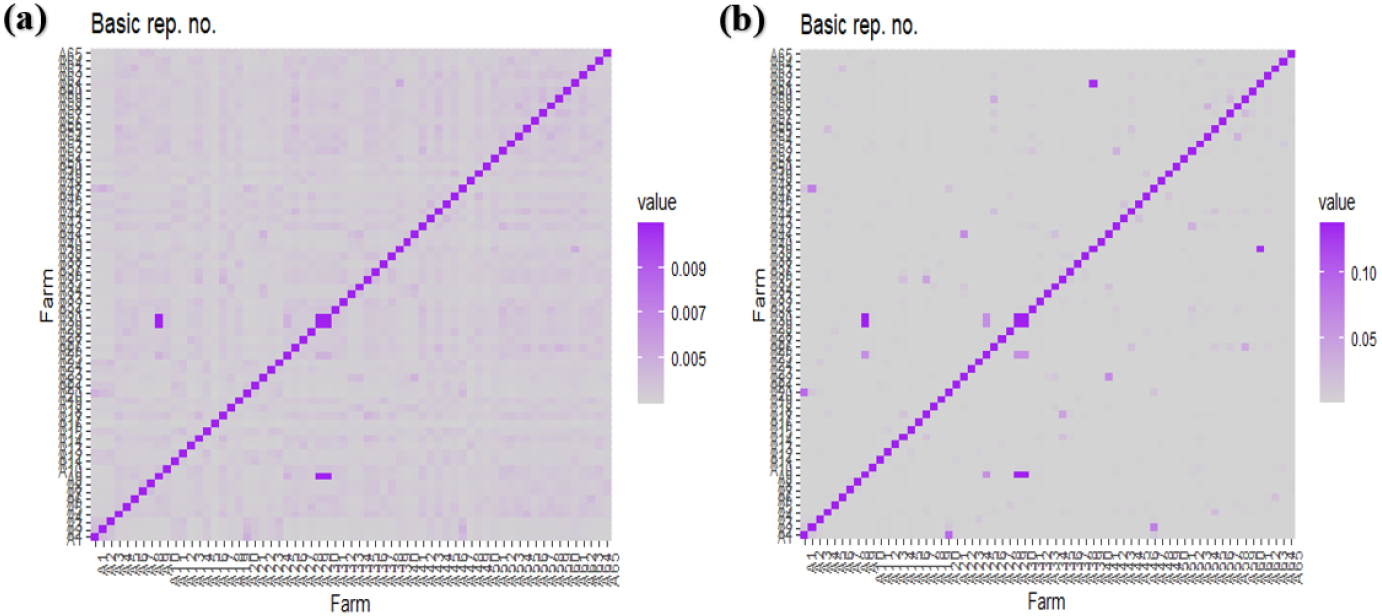
Estimated local reproduction numbers for the 2001 FMD outbreak in Mar Chiquita, using the fitted transmission kernel and infectious periods, before (a) and after (b) intervention.

**Figure A2:**
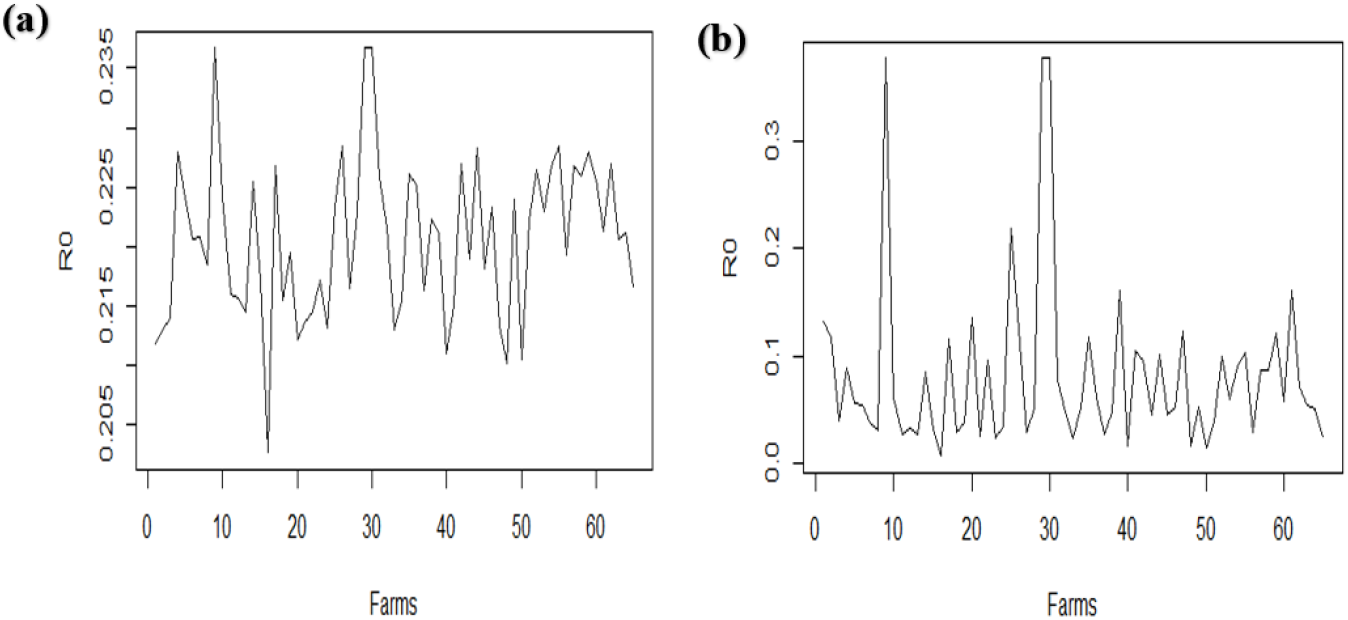
Simulated local reproduction number as a function of inter-farm distance, without (a) and with (b) intervention.

### A2 Daily transmission probability

To estimate the likelihood of daily outbreak progression, we computed the probability that a susceptible farm becomes infected on day *t* + 1, given the infection status on day *t*. This daily transmission probability is evaluated using the product of survival probabilities for all susceptible farms and infection probabilities for newly infected farms. The expression is given by:

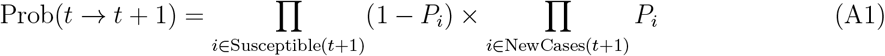

Here, *P*_*i*_ denotes the probability that farm *i* becomes infected on day *t* + 1, as computed from the spatial transmission model. This expression was used in the MCMC simulations to stochastically reconstruct transmission chains based on farm-level infection dynamics and spatial risk estimates.

